# Optimization of gene knockout approaches and practical solutions to sgRNA selection challenges in hPSCs with inducible Cas9 system

**DOI:** 10.1101/2024.07.09.602644

**Authors:** Jie Ni, Junhao Gong, Yuqing Ran, Rui Bai, Pufeng Huang, Zihe Zheng, Meng Zhou, Yi You, Feng Lan, Xujie Liu

## Abstract

**Rationale:** CRISPR/Cas9 has been extensively used to knock out genes, allowing the study of genetic loss-of-function in human pluripotent stem cells (hPSCs). However, the current use of the Cas9-sgRNA plasmid or iCas9 system for gene editing in hPSCs has resulted in limited and inconsistent editing efficiency, as well as labor-intensive work. Additionally, identifying single-guide RNAs (sgRNAs) with high cleavage efficiency and distinguishing them from ineffective ones, which efficiently induce frameshift INDELs (Indels and Deletions) but fail to eliminate target proteins expression, are major challenges in gene knockout experiments.

**Methods:** This study addresses above issues using an optimized doxycycline-induced spCas9-expressing hPSCs (hPSCs-iCas9) system. We initially developed this system by optimizing a number of parameters to maximize INDELs introducing efficiency in hPSCs-iCas9 cells. The INDELs determined by this system were then compared to predicted scores from three cleavage efficiency scoring algorithms to validate the algorithms’ accuracy and consistency. Furthermore, we conducted gene knockout using a set of sgRNAs targeting different exons of the ACE2 gene to achieve approximately 80% INDELs for each targeting locus. Western blotting was then performed to detect ACE2 protein expression levels, enabling the identification of potentially ineffective sgRNAs.

**Results:** Several critical factors, including cell tolerance to nucleofection stress, sgRNA stability, nucleofection frequency, and the cell-to-sgRNA ratio, were found to have significant impact on editing efficiency in hPSCs-iCas9. Fine-tuning these parameters markedly improved this efficiency, resulting in up to 93% INDELs for single gene knockout. The three scoring algorithms exhibited significant differences or even conflicts in scoring cleavage efficiency. Through comparing experimental observations to predicted scores, we discovered that the Benchling algorithm outperformed the other two in terms of accuracy and consistency. Furthermore, a sgRNA targeting exon 2 of ACE2 gene was quickly identified as ineffective, as evidenced by the edited cells pool containing 80% INDELs while ACE2 protein expression retained unchanged detected by Western blot.

**Conclusion:** The findings of this study offer valuable insights into the optimal design of gene knockout experiments in hPSCs and provide practical solutions to sgRNA selection challenges for gene editing.

## Introduction

Disease modeling using human pluripotent stem cells (hPSCs) and their derivatives has emerged as a pivotal approach for gene elucidation, mechanisms dissection, and drugs screening [1]. Within this field, gene knockout serves as a critical tool for investigating loss-of-function effects of disease-causing genes in hPSCs-based disease models. The CRISPR/Cas9 system, a revolutionary genome editing technology, has emerged as the preferred method for achieving precise and efficient gene knockouts. CRISPR/Cas9 functions by directing the Cas9 nuclease to a specific DNA sequence, resulting in double-strand breaks (DSBs). These DSBs are primarily repaired by the error-prone non-homologous end joining (NHEJ) pathway, often introducing insertions or deletions (INDELs) around the targeted site, thereby disrupting gene function.

The initial knockout efficiency of CRISPR/Cas9 in hPSCs was reported as low as 1-2% [2], primarily attributed to the inherent resistance of hPSCs to genome modification. Researchers have since devoted significant effort to exploring various variables that influence editing efficiency. These variables include nuclease delivery methods [3], sgRNA and homology arm design strategies [4–6], transfection techniques and targeted cells recovery approaches [7,8]. Additionally, improved Cas9 expression strategies have been developed, including Cas9 ribonucleoprotein (RNP) complexes [9,10], Cas9-modRNA [11] and inducible-Cas9. Among these, the doxycycline (Dox)-inducible spCas9 (iCas9) system stands out due to its cost-effectiveness, tunable nuclease expression, and remarkably enhanced INDELs efficiency, reaching up to 68% [12]. However, iCas9 has shown significantly variable targeting efficiency, which could range from 20% to 60% [12,13]. This variability is potentially attributed to a lack of comprehensive optimization in previous iCas9 studies.

In this work, we addressed this issue by optimizing a series of parameters within iCas9 system. These parameters included cell tolerance to nucleofection stress, transfection methods, sgRNA stability, nucleofection frequency, and the cell-to-sgRNA ratio. By systematically optimizing these factors, we were able to consistently achieve 82%-93% INDLEs efficiency in single gene knockout experiments. This represents a significant improvement in both efficiency and stability compared to previous iCas9 systems. We refer to it as the optimized hPSCs-iCas9 system.

Leveraging the exceedingly high gene knockout efficiency of the optimized hPSCs-iCas9 system, we aimed to broaden its applications beyond generating mutant hPSCs lines. Traditionally, designing sgRNAs for gene knockout often relies on online sgRNA design tools. These tools provide candidate sgRNAs along with predicted cleavage efficiency scores and off-target risks. However, this approach presents two significant challenges: 1) The cleavage efficiency scores predicted by the scoring tools are rarely validated by real experiments, raising concerns about their reliability; 2) The non-triplet reading frame shifts produced by some sgRNAs may fail to eliminate target protein expression, leading us to define such sgRNAs as ineffective in this study. This critical issue is often identified only after substantial time and effort have been invested in establishing the mutant hPSCs line. Therefore, how can we avoid selecting ineffective sgRNAs from the outset?

Here, to address the first concern, we experimentally determined the INDELs efficiencies of various sgRNAs by using the optimized hPSCs-iCas9 system, and then compared these experimental results to scores predicted by different online design tools. This comparison allowed us to identify the most reliable tool for in silico sgRNA design, considering both scoring accuracy and consistency with experimental results. Regarding the second issue, we propose a simple yet practical method utilizing Western blot to detect the target protein expression level in edited cells pools. Our hypothesis is that, within the context of our optimized system achieving high INDELs efficiency (reaching 80%), cells edited with effective sgRNAs will exhibit a significant reduction in target protein expression compared to those edited with ineffective sgRNAs. Western blotting readily detects these changes in protein levels, facilitating the identification of ineffective sgRNAs.

## Materials and Methods

### Cell culture and passage

Two ES cell lines, H9 (Wicell, WA09) and H7 (Wicell, WA07), two iPSC cell lines, U2 [14] and B1 [15], were cultured in PGM1 (Pluripotency Growth Master 1) Medium (Cellapy, Beijing, China) on Matrigel (Corning, USA) pre-coated multi-well plates at 37°C with 5% CO_2_ and passaged at 1:6 to 1:10 split ratio using 0.5 mM EDTA to dissociate cells when 80-90% cell confluency reached. Mycoplasma infection testing (Vazyme, Nanjing, China) was performed every other week to guarantee the cells were free of infection.

### hPSCs-iCas9 line construction

hPSCs-iCas9 line was created by inserting doxcyline-spCas9-puromycin cassette into the AAVS1 (also known as PPP1R12C) locus. In brief, two Addgene vectors #73500 and #113194 were co-electroporated into hPSCs at 1:1 weight ratio using 4D-Nucelofector (Lonza Bioscience, Germany) at program CA137. Addgene vector #73500 carries Tet-on^®^ 3G system and provides spCas9/anti-puromycin cassette flanked by homology arms of the AAVS1 locus. Vector #113194 delivers Cas9 and sgRNA targeting the AAVS1 locus. 48 hours after nucleofection, cells were selected with 0.5 to 1.5 μg/ml puromycin for one week, the surviving cell clones were subcloned, then genotyped by junction PCR [16] and Western blot. Primers and antibodies are indicated in supplementary Tables (Table S1 and S2). The pluripotency of the cells was validated by teratoma assay and detection of pluripotent hall markers expression.

### sgRNA, HDR donors design and synthesis

Unless otherwise specified, sgRNAs were designed by CCTop algorithms (https://cctop.cos.uni-heidelberg.de) [17] and listed in Table S3. sgRNA was either in vitro transcribed (IVT-sgRNA) using EnGen® sgRNA Synthesis Kit (New England Biolabs, USA) or chemical synthesized and modified (CSM-sgRNA) by GenScript Corporation (Nanjing, China). The CMS-sgRNA harbors 2’-O-methyl-3’-thiophosphonoacetate in both 5’ and 3’ ends to enhance sgRNA stability within cells [5]. CCTop was also used to search potential off-target sites. The top ten sites for TAZ gene targeting sgRNA were checked by PCR Sanger sequencing (Table S4).

For point mutation knock-in study, we selected L275F mutation in the C1QBP gene [18] as our target. Introducing this mutation will eliminate a PAM (AGG>AAG on the antisense strand), naturally preventing Cas9-sgRNA complex from binding to and recutting the mutated allele (Figure S2C). We utilized single-stranded oligodeoxynucleotides (ssODNs) as the HDR (homology directed repair) donor template and designed a 100-nucleotides (nt) ssODN complementary to the sgRNA strand [2,8], featuring homology arms flanking the mutation site with symmetric extension. The ssODNs were synthesized by Sangong Biotech (Shanghai, China) and listed in Table S3.

### Nucleofection of sgRNA and ssODN

For Dox-treated hPSCs-iCas9 nucleofection, cells were dissociated with EDTA and pelleted by centrifugation at 250g for 5 minutes. sgRNA or sgRNA/ssODN mix was combined with nucleofection buffer (P3 Primary Cell 4D-Nucleofector™ X Kit) and electroporated into cell pellets using the CA137 program on Lozon Nucleofector. Repeated nucleofection was conducted 3 days after the first nucleofection following the same procedure. To target multiple loci, we conducted nucleofection with two or three sgRNAs at the same weight ratio to a fixed amount of 5 μg.

### hPSCs-CMs differentiation and phenotypes characterization

Monolayer hPSCs differentiated cardiomyocytes (hPSCs-CMs) were induced following the previous protocol [19] with minor modifications. Briefly, hPSCs were detached by incubating with 0.5 mM EDTA for 5 min before being seeded at a density of 25,000 cells/cm^2^ into Matrigel pre-coated 6-well plates. Upon reaching 90% confluency, cells were treated with cardiomyocytes differentiation basic medium (2 mM L-glutamine RPMI-1640 plus 1x B27 minus insulin) containing 5 μM CHIR99021 (STEMCELL Technologies, Canada) for 48 hours, followed by basic medium containing 5 μM IWR-1-endo (STEMCELL Technologies) for another 48 h. The cells were then cultured in basic medium for more than 6 days until spontaneous contractions were observed. The hPSCs-CMs were then enriched by lactate selection medium [20] for 3 days before being replaced with basic medium for long-term culture. hPSCs-CMs were used for research between 30 and 40 days after differentiation. The purity of hPSCs-CMs was determined using flow cytometry and immunofluorescence to detect the expression of the cardiomyocyte-specific sarcomeric protein cTNT (Cardiac Troponin T). To visualize mitochondria, hPSCs-CMs were incubated with 25 nM Mitotracker Green (Life Technologies, USA) in PBS for 15 min at room temperature, followed by three washes prior to confocal imaging. Furthermore, the Mito stress assay (Seahorse, Agilent, USA) was conducted to evaluate mitochondrial dysfunctions in TAZ gene knockout (TAZ-KO) hPSCs-CMs.

### Edits analysis algorithms validation

Unless otherwise states, Sanger Sequencing chromatograms were analyzed using ICE (Inference of CRISPR Edits) [21] (https://ice.synthego.com/#/) in this study. To validate the sensitivity and accuracy of this algorithm, we compared it to another algorithm TIDE (Tracking of Indels by Decomposition) [22] (https://tide.nki.nl), and T7 endonuclease I (T7EI) mismatch assay (Beyotime Biotechnology, Shanghai, China) [8,12]. Leveraging our findings that cell tolerance to nucleofection and cell-to-sgRNA ratio influence editing efficiency, we generated three distinct edited cells pools with progressively increasing INDELs levels. These pools were created by nucleofecting hPSCs-iCas9 cells with varying parameters, 1#: 1 μg sgRNA for 4×10⁵ H7-Cas9 cells, 2#: 5 μg sgRNA for 8×10⁵ H7-Cas9 cells, and 3#: 5 μg sgRNA for 8×10⁵ H9-Cas9 cells. Genomic DNA was extracted from these cell pools for PCR amplification. The PCR products were then subjected to either T7EI assay or Sanger sequencing, followed by ICE and TIDE analyzing. ImageJ software was employed to measure gray values of PCR products digested by T7EI and the INDELs percentage was calculated as previously reported [12].

Next, we validated ICE analysis against actual edits of single-cell sorted clones. In brief, a sgRNA targeting the PLA2G6 gene was electroporated into Dox-treated H9-iCas9 cells. After 3 days, half electroporated cells were harvested for genomic DNA extraction, PCR amplification, Sanger sequencing and ICE analysis. The remaining half cells were subjected to single-cell sorting and deposited into a 96-well plate at one cell per well using the CytoFLEX SRT Cell Sorter (Beckman, USA). After approximately 10 days of culture, the actual editing spectrum was calculated from observed genotyping results of 50 sorted cell clones.

### Protein expression detection

Protein was extracted from RIPA buffer lysed cells supplemented with protease inhibitor cocktail (Roche, Basel, Switzerland). After quantification by Pierce BCA protein assay kit (Life Technologies, USA), protein extracts were diluted to 1.0 μg/μl for Western-blot. For the indirect immunofluorescence assay (IFA), cells were fixed in 4% PFA and 0.1% triton X-100 in PBS for 60 min at room temperature. After three washes, cells were incubated with primary antibodies overnight at 4℃. The following day, cells were washed three times before being incubated with secondary antibodies and DAPI at room temperature for 1.5 h prior to confocal imaging. Primary, secondary antibodies information and their optimal dilution are indicated in Table S3.

### Statistical analysis

Unpaired two-tailed t-tests were conducted using GraphPad Prism software to compare two groups. The data are presented as means ± SD unless otherwise specified. P values ≥ 0.05 were considered non-significant; significance levels were defined as *P < 0.05, **P < 0.01, ***P < 0.001, and ****P < 0.0001. Statistics were performed with GraphPad Prism 8.

## Results

### Multiple challenges associated with Cas9-sgRNA plasmid approach

A typical protocol for creating mutant hPSCs lines using the Cas9-sgRNA plasmid comprises four steps: plasmid transfection, antibiotic selection, cells sub-cloning, and clonally expansion (Figure 1A). We tested transfection efficiency by nucleofecting four hPSCs lines with a Cas9-sgRNA vector (PX458-T2, Addgene #113194) targeting the AAVS1 locus at manufacturer’s recommended cell-to-sgRNA ratio (5 µg plasmid per 0.8 million cells). This vector contains a GFP marker for flow cytometry testing of transfection efficiency. With the exception of B1, which achieved 16.81% transfection, the other three lines had efficiencies below 6%, with H9 cells having the lowest at 2.39% (Figure 1B). Even after doubling the plasmids amount to 10 µg, the efficiency remained below 8% (Figure 1C). Meanwhile, overloaded plasmids caused obvious cytotoxicity and distorted cells morphology (data not shown).

**Figure 1.**
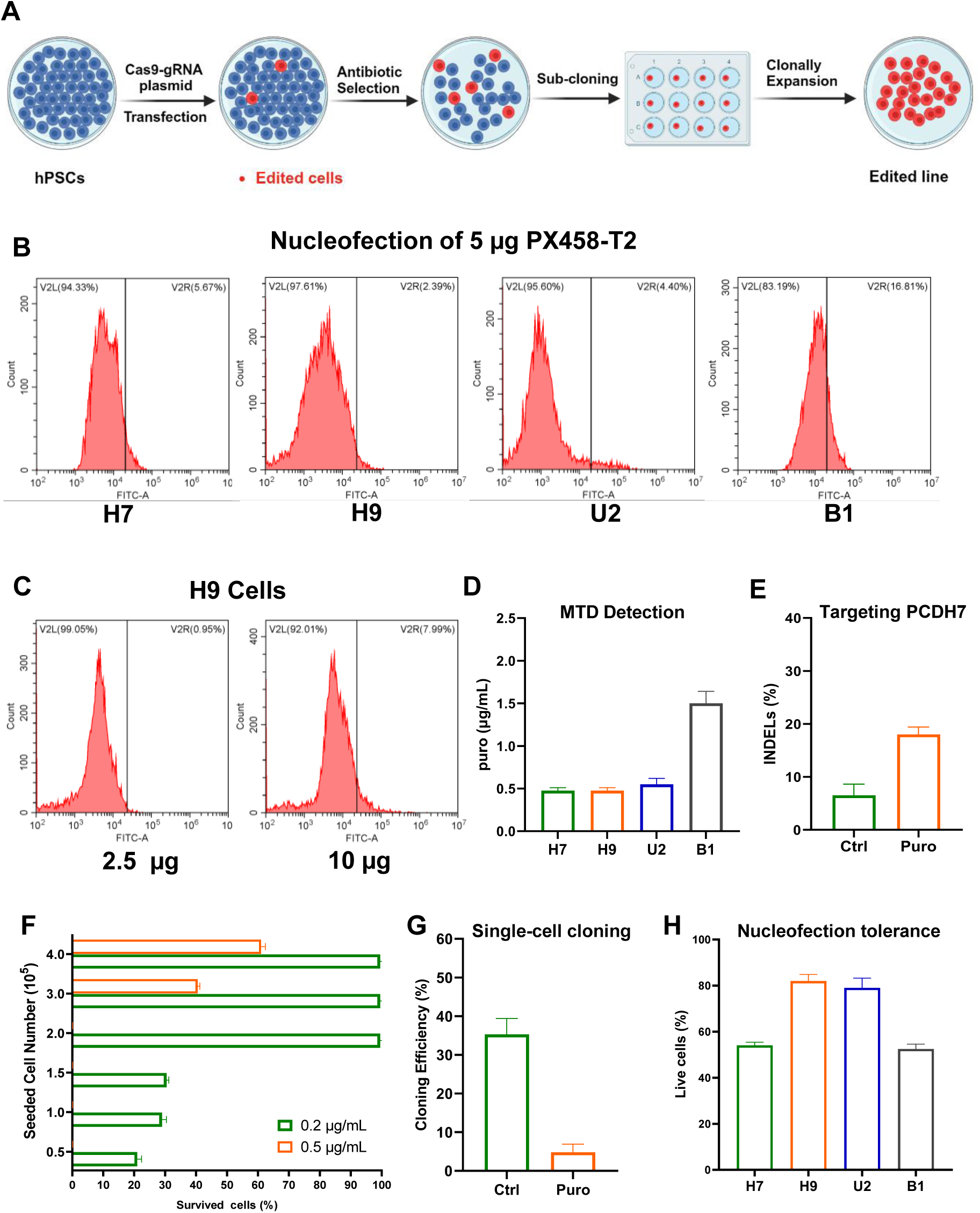
Limitation s of Cas9-sgRNA plasmid for gene knock in hPSCs. (A) Conventional workflow for constructing a mutant cell line using Cas9-sgRNA plasmid. (B) Flow cytometry analysis of the GFP-positive cells after various hPSCs lines nucleofected with 5 μg PX458-T2 plasmids. (C) FACS analysis of the GFP-positive cells H9 line following H9 cells nucleofected with either 2 μg or 10 μg of PX458-T2 plasmids. (D) Determination of the maximum tolerated dose (MTD) of puromycin for H9 cells measured at 30% cell confluency. (E) INDELs enrichment efficacy by puromycin selection. (F) The impact of H9 cell confluency on puromycin selection efficacy. Data were normalized to the mock treatment controls. (G) Quantitative analysis of cloning efficiency before and after puromycin selection. (H) Quantitative assessment of cell viability at 24 hours post-nucleofection. Data were normalized to the mock treatment controls. All experiments were performed in triplicate (n=3). Error bars in the graphs represent the mean ± standard deviation (SD).

Antibiotic positive selection is commonly employed to enrich target cells, here we ran a series of tests to gauge its consistency and sensitivity. When puromycin selection was initiated at 30% cell confluency, B1 cells exhibited exceptional resistance to puromycin, with maximum tolerated dose (MTD) of 1.5 µg/ml, while rest lines were highly sensitive to puromycin, showing an average MTD of 0.5 µg/ml (Figure. 1D). This suggests that innate antibiotics sensitivity varies across hPSCs lines, emphasizing the necessity of determining each line’s MTD before selection work. Next, the selection efficacy was evaluated by nucleofecting H9 cells with a puromycin-resistant Cas9-sgRNA plasmid targeting the PCDH7 gene, followed by 5 days of puromycin selection at concentrations of 0.5-1.0 µg/ml. Upon examining INDELs enrichment, we found that the selecting above MTD only elevated INDELs from 5% to 18% (Figure. 1E). This indicates that antibiotic selection is less effective than initially anticipated. One contributing factors to this ineffectiveness is the cell confluence at the start of selection. When H9 cells reached 80% confluence (4×10^5^ cells seeded per well in 12-well plate) to start selection, the MTD of 0.5 μg/ml determined from 30% confluence was only capable of killing fewer than half cells. While 0.2 μg/ml puromycin, which effectively eliminated 80% of cells for starting with 0.5×10^5^ million seeding cells, entirely lost its selection capacity for 2×10^5^ million seeding cells (Figure. 1F). We further investigated the impact of antibiotic selection on single-cell cloning efficiency. Cells not treated with puromycin exhibited an average cloning efficiency of 35%. In contrast, those subjected to puromycin treatment showed a significantly lower efficiency, dropping below 5% (Figure. 1G). This finding suggests that excessive stress imposed by antibiotic selection renders cells more susceptible to single-cell manipulation.

In summary, the experimental data presented above underscore the challenges associated with the Cas9-sgRNA plasmids approach, highlighting the critical need to establish a simple, efficient and consistent system for effective gene editing in hPSCs.

### hPSCs-iCas9 lines generation and characterization

During the nucleofection experiments on various hPSCs lines, we observed that these lines exhibited varying levels of tolerance to nucleofection. The robust lines: H9 and U2, retained more than 80% viability following nucleofection. In contrast, the vulnerable lines: H7 and B1 had a viability of approximately 50% (Figure. 1H). Considering that significant stress caused by nucleofection may potentially influence gene editing efficiency, we selected two different stress-tolerant lines H9 and H7 for constructing the hPSCs-iCas9 lines (designated as H9-iCas9 and H7-iCas9, respectively) (Figure 2A) and for subsequent experiments. Further investigation revealed that these two hPSCs-iCas9 lines retained the ability to differentiate into tissue derivatives of the three embryonic germ layers in teratoma assays (Figure S1A), and expressed high level of pluripotency markers OCT4 and NANOG (Figure S1B). Furthermore, karyotyping analysis confirmed that targeting AAVS1 did not cause detectable chromosomal aberrations (Figure S1C).

**Figure 2.**
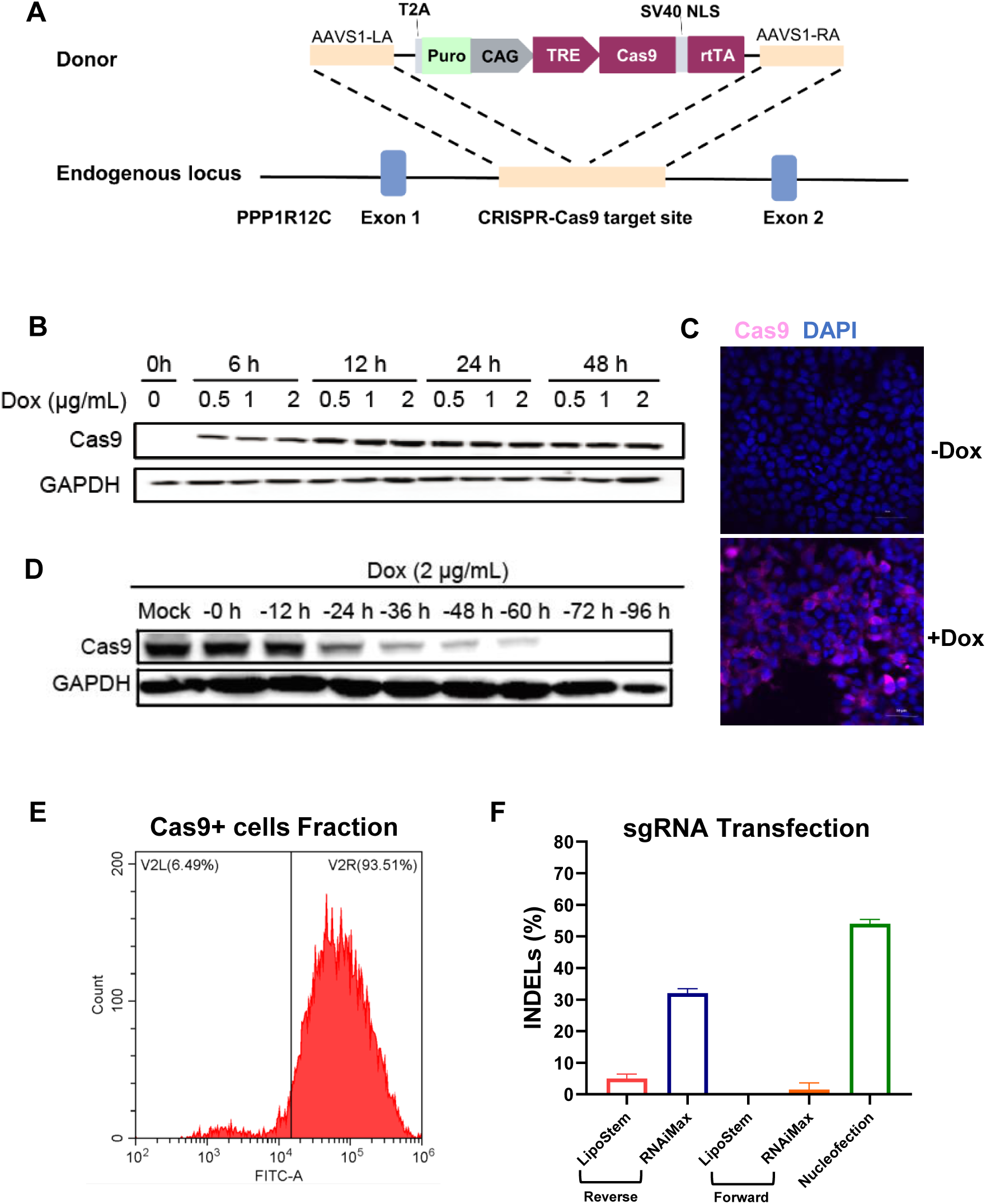
Generation and characterization of hPSCs-iCas9. (A) Generation of hPSCs-iCas9 lines through CRISPR/Cas9-mediated gene targeting at the AAVS1 locus. (B) Determination of the optimal conditions for Cas9 introduction in H7-iCas9 cells. (C) Immunofluorescence assay (IFA) detection of Cas9 protein expression in H7-iCas9 after 12 hours of treatment with 0.5 μg/ml Dox. (D) Western blot analysis of Cas9 expression persistence following Dox removal in H7-iCas9 cells, with Dox not removed in mock cells. (E) Flow cytometry analysis of the fraction of cells expressing Cas9 after 12 hours of treatment with 0.5 μg/ml Dox. (F) INDELs efficiency achieved using different transfection methods targeting the TAZ gene. Data represent mean ± SD, n=3.

Western blotting was then conducted to determine the optimal conditions for Cas9 induction. Treating hPSCs-iCas9 with 0.5 μg/ml Dox for 12 hours was sufficient to induce a high level of Cas9 expression (Figure 2B, C and E). This expression persisted strongly for the first 12 hours following Dox withdrawal, sharply declined at 24 hours, and became undetectable by 72 hours (Figure 2D), suggesting the optimal time window for gene editing is within the first 12 hours after Dox removal.

Although Cas9 protein was not detectable by Western blot in the absence of Dox, nuclease leakage remains a safety concern in Tet-On system. To investigate potential leakage, we performed nucleofection of H9-iCas9 with sgRNAs targeting the TAZ or SCN5A gene in the absence of Dox. The subsequent Sanger sequencing analysis by ICE revealed no detectable edits in both genes (Figure S1D).

### Efficient transfection methods determination

Given that the Cas9 nuclease is ubiquitously inducible upon Dox addition, in hPSCs-iCas9 cells, the only requirement for achieving genome knockout is efficient delivery of sgRNA. Here, we compared three widely used methods/reagents: nucleofection, LipoStem, and RNAiMAX to assess their efficacy in delivering CMS-sgRNA targeting the TAZ gene. As expected, Nucleofection yielded an average of 55% INDELs efficiency in Dox treated H9-iCas9 cells, significantly higher than LipoStem and RNAiMax, whether used in forward or reverse transfection (Figure 2F). Thus, we proceeded with nucleofection for parameters optimization in subsequent exploration. Intriguingly, LipoStem exhibited highest efficiency with remarkable 91.09% GFP-positive rate in GFP-mRNA transfection experiments, while Nucleofection came in second with 67.37% (Figure S1E). We hypothesize that the disparities in data observed between gRNA and mRNA are due to the different cellular compartments they need to reach to function. gRNA must enter the cell nucleus to function, while GFP-mRNA only needs to be translated in the cytoplasm. Consequently, nucleofection, which is promoted as a method to deliver molecules directly into the cell nucleus, achieves higher editing efficiency.

### Parameters optimization to maximize INDELs efficiency

Having determined the transfection method for hPSCs gene editing, we proceeded to investigate the parameters that potentially affect INDELs efficiency. We initially assessed the performance of IVT-sgRNA and CSM-sgRNA in both H7-iCas9 and H9-iCas9 cells. As expected, CSM-sgRNA, which benefits from enhanced stability through chemical modification, exhibited superior performance over IVT-sgRNA in both hPSCs-iCas9 lines. INDEL efficiency achieved with CSM-sgRNA was observed to be up to three-fold and two-fold greater than IVT-sgRNAs in H7-iCas9 and H9-iCas9 cells, respectively (Figure 3A). More importantly, editing efficiencies were consistently higher in robust H9-iCas9 cells compared to the more vulnerable H7-iCas9 cells, regardless of sgRNA types (CMS or IVT). This confirmed that cells’ tolerance to nucleofection stress plays a crucial role in editing efficiency. Additionally, we unexpectedly observed that the cells harvest time affects the efficiency measurement. In CMS-sgRNA experiments, INDELs efficiency was lower during the first two days after nucleofection, gradually increasing and stabilizing by the third day (Figure 3A). However, following its peak on day 3, a gradual and sustained decline in INDELs efficiency was observed, with values dropping from 80% to 15% between day 3 and day 60 post-nucleofection. (Figure S2A). One potential explanation to this observation could be that efficient genome editing mediated DSBs activates the p53 pathway, causing programmed cell death in the edited cells [23]. Based on these findings, we proceeded with subsequent experiments using CSM-sgRNA and conducted analysis on the third day of post-nucleofection.

**Figure 3.**
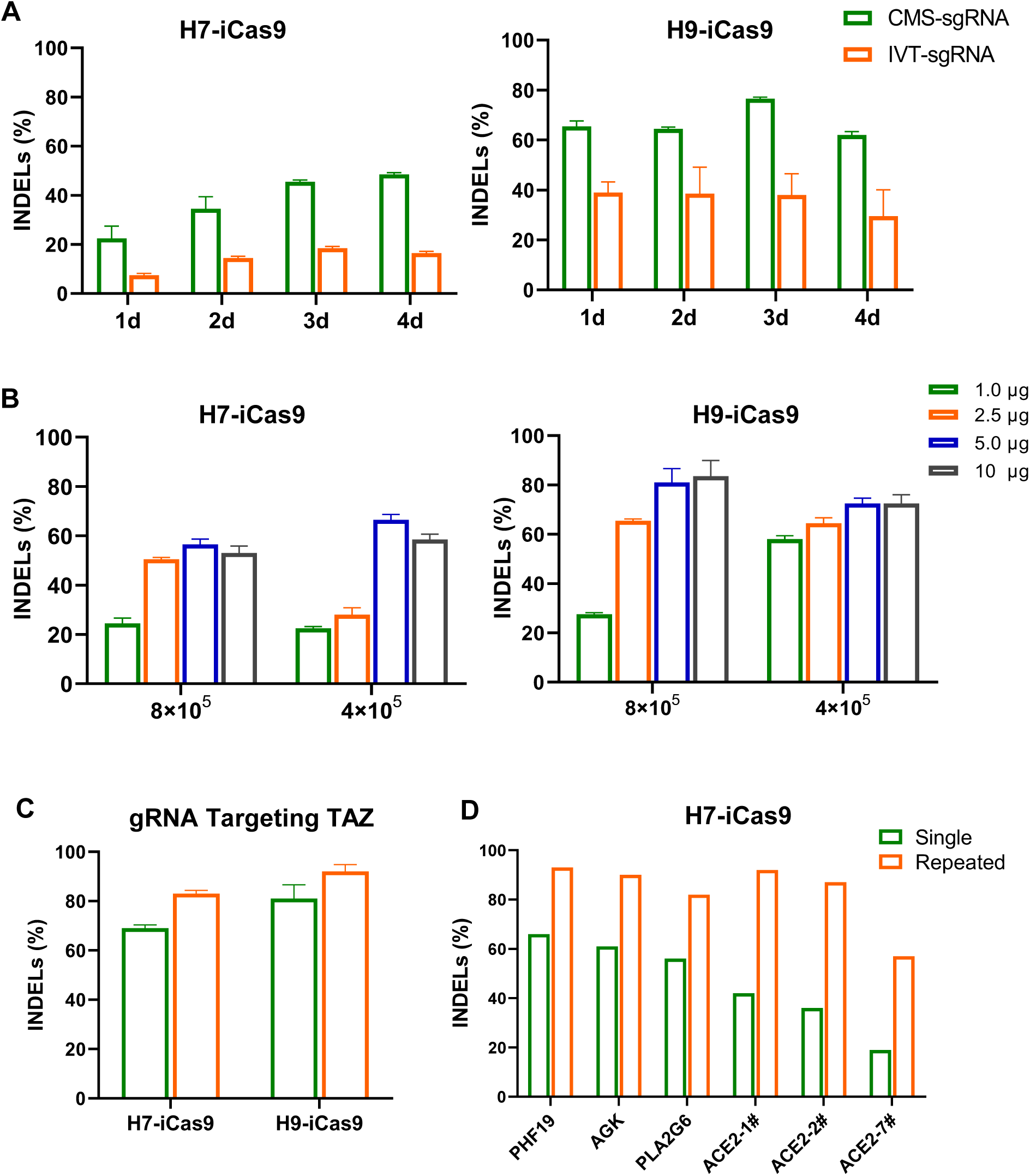
Parameter optimization for enhancing gene knockout efficiency. (A) Comparison of INDELs introducing efficiency between CMS-sgRNA and IVT-sgRNA. Nucleofected cells were analyzed from 1 to 4 days post-nucleofection (B) Investigation of the optimal cell-to-sgRNA ratio to enhance INDELs efficiency. (C) and (D) repeated nucleofection (Repeated) significantly increased the INDELs efficiency compared to single time nucleofection (Single) using various sgRNAs targeting different genes. Experiments in (A) to (C) utilized targeting TAZ gene sgRNA, performed in triplicate (n=3). Data represent mean ± SD. N=1 in (D).

We next investigated the role of cell-to-sgRNA ratio in gene editing efficiency. By nucleofection of increasing sgRNA (targeting the TAZ gene) amount from 1.0 to 10 μg into either 4×10^5^ cells using Nucleocuvette® Strip or 8×10^5^ cells using single Nucleocuvette®. We discovered that 5 μg of sgRNA for 4×10^5^ H7-iCas9 cells and 10 μg of sgRNA for 8×10^5^ H9-iCas9 cells yielded maximum INDELs efficiency, averaging 66.5% and 83.5%, respectively (Figure 3B). Different to plasmids, excessive sgRNA did not induce apparent cytotoxicity to cells (data not shown). Since 5 μg sgRNA produced similar INDELs efficiency as 10 μg (81% vs. 83.5%) in 8×10^5^ H9-iCas9, and considering the costs-effectiveness, we opted for 5 μg of sgRNA for 8×10^5^ H9-iCas9 and 4×10^5^ H7-iCas9 cells in subsequent experiments.

Next, we performed consecutive two times nucleofection (repeated nucleofection) of sgRNAs and surprisingly found that repeated nucleofection substantially contributes to improving INDELs efficiency. Firstly, repeated nucleofection further enhanced the already high INDELs efficiency, increasing it from 69% to 83% in H7-iCas9 cells and from 81% to 93% in H9-iCas9 cells (Figure 3C). Secondly, repeated nucleofection dramatically improved the initial lower INDELs efficiency, for instance, raising it from 36% to 87% in H7-iCas9 using ACE2-2# sgRNA (Figure 3D). Lastly, by incorporating repeated nucleofection, the optimized Cas9 system consistently achieved INDELs efficiencies of 82-93% in both robust and vulnerable hPSCs-iCas9 lines (Figure 3C, D). We also found that the effectiveness of repeated nucleofection in enhancing INDELs has a limit, which is threefold. This is evidenced by experimental data showing that using ACE2-7# sgRNA increased INDEL efficiency from 19% to 57%. In other words, sgRNA’s intrinsic cleavage efficiency sets the ceiling for parameters optimization. Thus, we recommend prioritizing sgRNAs that achieve at least 30% INDEL efficiency from the first nucleofection for subsequent experiments, as this threshold ensures that the final efficiency can be effectively boosted to over 80% through repeated nucleofection.

### Efficient multiplexed gene knockout and DNA fragment deletion

Through step-by-step optimization, we have developed the optimized hPSCs-iCas9 system for efficient gene knockout at single loci. We next examined the performance of this system in multiple locus knockout and DNA fragment deletion. We started with dual genes knockout by co-nucleofecting two sgRNAs targeting the TAZ and SCN5A gene. Through repeated nucleofection, the SCN5A and TAZ gene obtained an average of 88% and 81% INDELs, respectively (Figure 4A). Also, the optimized system demonstrated remarkable performance in triple gene knockout study, with the TAZ, PHF19 and ACE2 gene obtaining an average of 78%, 76% and 80% INDELs, respectively (Figure 4B).

**Figure 4.**
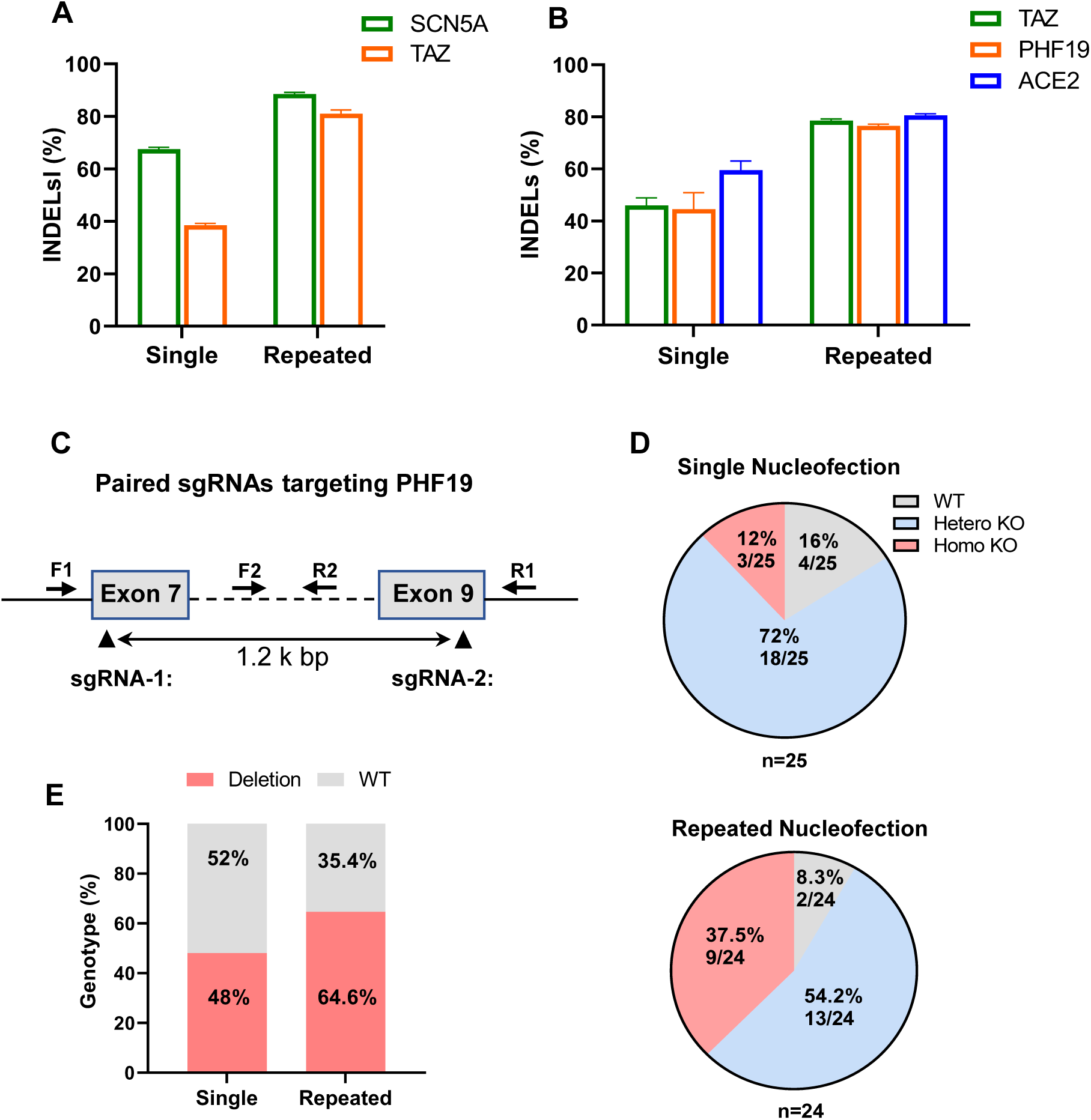
Efficient editing in multiplexed knockout and DNA fragment deletion. (A) and (B) The optimized H9-iCas9 showed highly efficiency in double or triple genes knockout experiments, respectively. Data are shown as mean ± SD, n=3. (C) Schematic illustration of genotyping primers design and paired sgRNAs spanning exon 7 to 9 on the PHF19 gene. (D) Genotypes of single-cell sorted cell clones from single and repeated nucleofection. Pie charts illustrate the fractions of homozygous, heterozygous gene knockout, and wild-type cell clones. (E) Quantitative analysis of mutant and wild-type allele fractions from single and repeated nucleofection.

In addition to NHEJ-mediated small INDELs, the strategy of removing large DNA fragments spanning multiple exons is commonly adopted to knockout genes. Here, we designed a pair of sgRNAs (paired sgRNAs) spanning exon 7 to exon 9, covering a 1.2 kb region of the PHF19 gene (Figure 4C). After single and repeated nucleofection, cells were single-cell sorted and deposited into 96-well plates. Following about 10 days of culturing, single-cell derived cell colonies were collected for PCR genotyping using primers F1, R1 and F2, R2 to identify WT, homozygous (biallelic) and heterozygous (monoallelic) knockout genotypes (Figure S2B). Compared to 12% (3/25) homozygous and 72% (18/25) heterozygous knockout clones obtained from single nucleofection, repeated nucleofection significantly increased homozygous knockout fraction to 37.5% (9/24) and reduced heterozygous knockout to 54.2% (13/24), simultaneously decreased wild-type fraction from 16% (4/25) to 8.3% (2/24) (Figure 4D). In terms of edited alleles proportion, single nucleofection generated excision to 48% alleles, while repeated nucleofection increased it to more than 64% (Figure 4E).

In addition to knockout, point mutation knock-in via HDR was also tested in the optimized system. By varying the ratio of sgRNA to ssODN in H9-iCas9, we yielded an average efficiency of 18% in single nucleofection and 25% in following repeated nucleofection at 1:1 sgRNA-to-ssODN ratio (Figure S2D).

### Cleavage efficiency scoring validation

To assess the accuracy of sgRNA cleavage efficiency scoring algorithms, we designed five sgRNAs (TAZ-1# to TAZ-5#, Table S3) targeting the exon 6 of the TAZ gene and scored them using three online algorithms: CCTOP, Deephf and Benchling. Significant differences were observed among the predicted scores from these algorithms (Figure 5A). For instance, 3# sgRNA received a moderate score from CCTOP (0.77) and Benchling (57.4), but markedly low score from DeepHF (0.005). Conversely, 4# sgRNA received the highest score from CCTOP (0.85) and DeepHF (0.42), but the lowest score form Benchling (33.3). To validate these predictions, we determined the INDELs efficiency of these 5 sgRNAs using the optimized hPSCs-iCas9 system with single nucleofection. Sanger sequencing revealed that 3# sgRNA efficiently introduced INDELs with an average efficiency of 64%, Conversely, sgRNA 4# was the most inefficient in this regard, yielding an average efficiency of only 23%. These data suggests that Benchling is superior to the other two algorithms in terms of accuracy and consistency for predicting sgRNA efficiency. This finding was further corroborated by experiments evaluating sgRNAs targeting different genes (Figure S2E). For instance, sgRNA 2# targeting PLA2G6 received the lowest score (0.51) from CCTOP, a second-highest score (0.56) from Deephf, and the highest score from Benchling (72.4). Consistent with Benchling’s prediction, experimental results confirmed sgRNA 2# to be the most efficient in introducing INDELs. Pearson correlation analysis further supported our findings, demonstrating the strongest correlation (R^2^=0.4943) between Benchling’s computational predictions and the experimental results, Conversely, Deephf exhibited the minimal correlation (R^2^=0.0005), while CCTOP even showed a negative correlation (Figure 5B).

**Figure 5:**
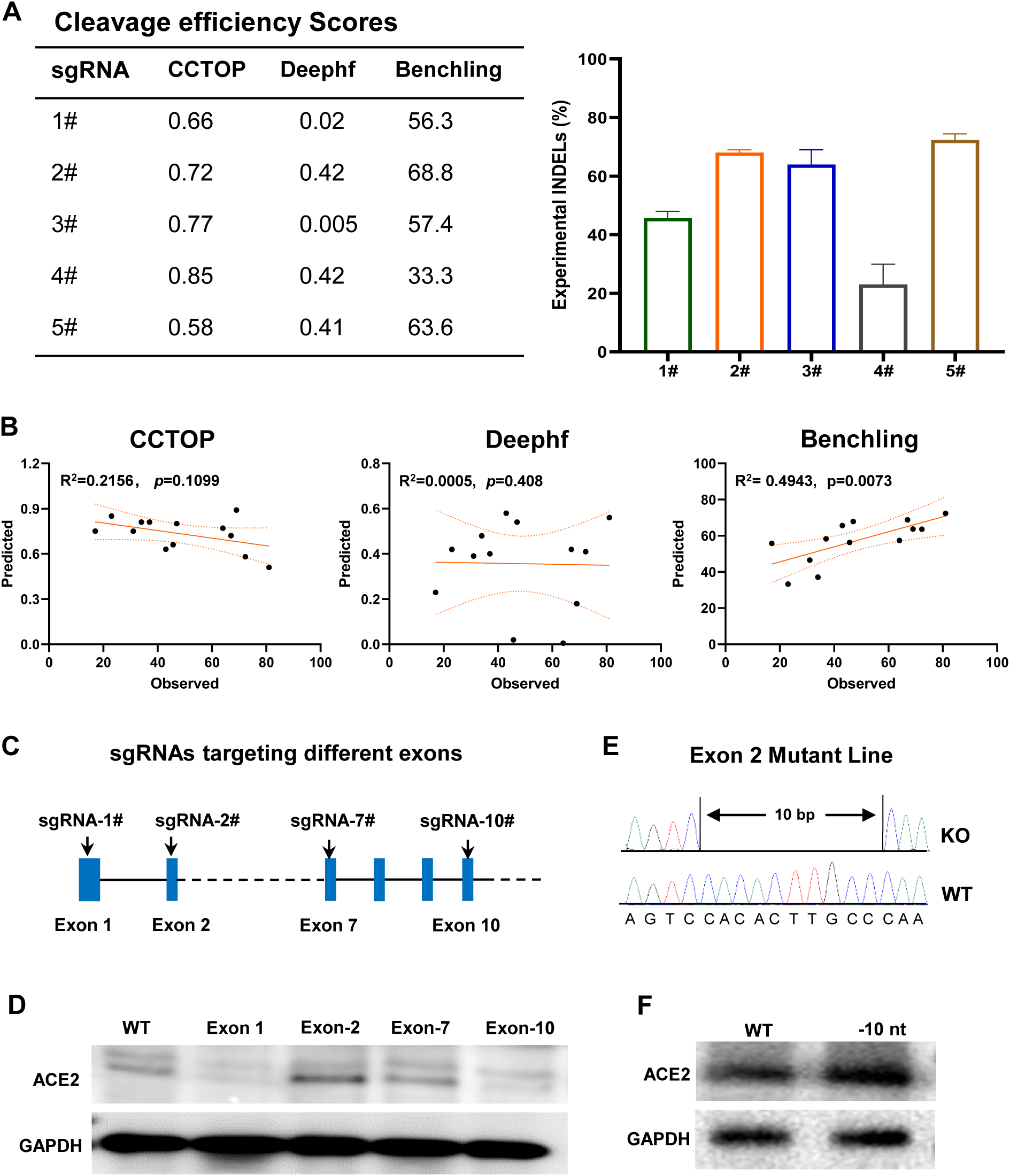
Identification of cleavage-efficient and ineffective sgRNA. (A) Validation of cleavage efficiency scores against the actual edits. Left: Cleavage efficiency predicted by three different scoring algorithms. Right: Experimental INDELs efficiency of sgRNAs (1# to 5#) targeting the TAZ gene determined in H9-iCas9. Data are shown as mean ± SD, n=3. (B) Scatter plots with regression showing the correlation between observed INDELs efficiency and predicted cleavage efficiency scores from three scoring algorithms (95% CI). (C) Schematic illustration of sgRNAs design targeting different exons of the PHF19 gene. (D) Western blot analysis of ACE2 expression with sgRNAs targeting different exons of the ACE2 gene. (E) The genotype of a cell line established with a biallelic 10 nt deletion (-10 nt) in exon 2 of the ACE2 gene. (F) Western blot analysis shows the ACE2 protein is not eliminated in cell line with the 10 nt deletion in exon 2.

### Identification of ineffective sgRNA

We designed four sgRNAs targeting exon 1, 2, 7, and 10 of the ACE2 gene, respectively (Figure 5C). After repeated nucleofection, INDELs efficiency reached 87%, 87%, 58%, and 79% for each exon, respectively (Figure S3A). Western blot analysis revealed that cells edited with sgRNA targeting exon 1 and 10 displayed a significant reduction in ACE2 expression, while targeting exon 7 showed no substantial change, and targeting exon 2 even exhibited increased expression (Figure 5D). Based on this observation, we suspected that sgRNA targeting exon 2 was the ineffective sgRNA, while those targeting exons 1 and 10 were effective. For the sgRNA targeting exon 7, which only achieved moderate INDELs efficiency, likely suffered from intrinsically low cleavage efficiency (Figure 3D), making it difficult to determine its effectiveness. To validate our speculation, we conducted single-cell sorting to recover edited cells and eventually established a cell line with a biallelic 10 nt deletion in exon 2 (Figure 5E). Subsequent Western blot showed that ACE2 protein expression was not abolished in this cell line (Figure 5F), confirming this sgRNA is ineffective.

### Characterization and gene editing of hPSCs-iCas9-CMs

The final step in generating an hPSCs disease model involves differentiating the mutation-carrying hPSCs-iCas9 into the desired functional cells. Our data indicated that the integration of iCas9 cassette had little impact on cardiomyocyte induction, both H7 and H7-iCas9 cells could be efficiently differentiated into cardiomyocytes, with up to 90% purity determined by cardiac sarcomeric protein cTNT expression (Figure 6A, B). Following this, we investigated the disease modeling capabilities of hPSCs-iCas9 by evaluating the disease phenotypes of TAZ knockout H7-iCas9-CMs. TAZ deficiency is known to disrupt cardiolipin remodeling, leading to severe mitochondrial dysfunction and ultimately causing Barth Syndrome [24]. We created a cell line harboring a homozygous 2 nt deletion using the optimized hPSCs-iCas9 system and designated it as TAZ-KO-H7-iCas9 (Figure 6C). Upon differentiation into cardiomyocyte, these cells exhibited a dramatic loss of mitochondrial membrane potential and activity (detected by Mito-Tracker, Figure S3B), accompanied by severely impaired mitochondrial function as assessed in a Mito stress assay (e.g., maximum respiration capacity and ATP production) (Figure 6D, E).

**Figure 6.**
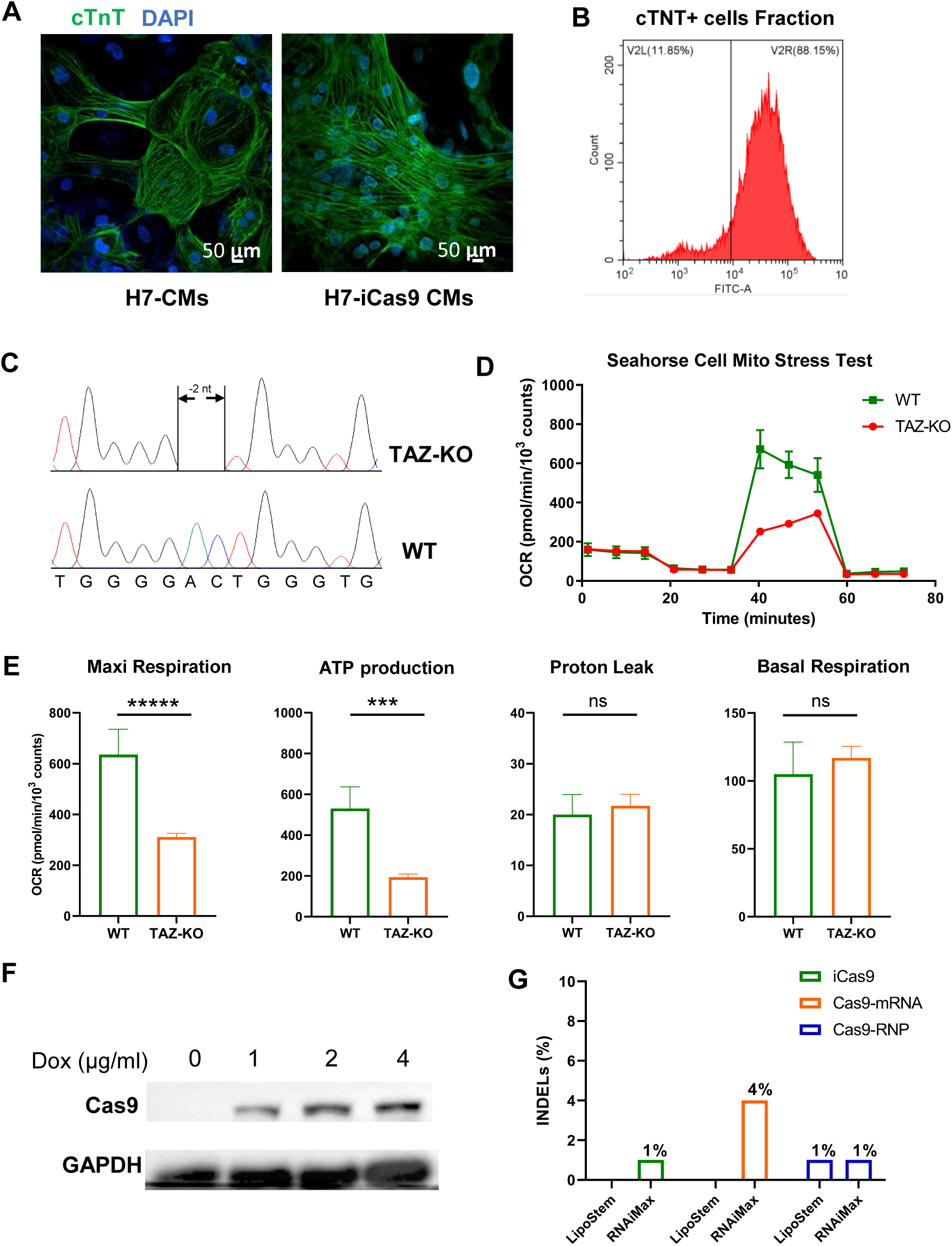
hPSC-CMs Phenotypic characterization and gene editing. (A) Representative images of cTNT expression in H7-CMs and H7-iCas9-CMs. Cells are counterstained with DAPI to visualize cell nucleus. Scale bar=50μm. (B) Flow cytometry analysis of the proportion of cTNT-positive cells, indicating the high purity (>88%) of differentiated cells. (C) Establishment of a TAZ gene knockout cell line with a biallelic 2 nt deletion in exon 6. (D) and (E) Seahorse assay detection of mitochondrial dysfunction in TAZ knockout hPSC-CMs. (D) Representative traces of the Mito stress test. Oxygen consumption rate (OCR) was normalized to viable cell counts. (E) Statistical analysis of basal OCR, OCR attributable to ATP production, proton leak, and maximal respiratory capacity. n=5. Data are shown as mean ± SD, Two-tailed t-test: ***P<0.001, *****P<0.0001. (F) Exploration of Cas9 protein introduction conditions in H7-iCas9-CMs through Western blot analysis. (G) Performance of iCas9, Cas9-RNP and Cas9-mRNA in introducing INDELs using liposomal delivery reagents.

We also attempted to perform gene editing on Dox-treated H7-iCas9-CMs. With Dox treatment for 48 h, Cas9 was successfully induced at the concentration of 1 μg/ml (Figure 6F). Since nucleofection buffers and programs specifically optimized for hPSCs-CMs were unavailable, we explored alternative delivery methods using liposome transfection reagents (LipoStem and RNAiMAX). However, these methods resulted in consistently low INDELs efficiencies (Figure 6G). We also tested RNP-Cas9 and mRNA-Cas9 delivery, but these approaches yielded similarly low efficiencies. These findings suggest that liposome-based transfection is not suitable for gene editing in hPSCs-CMs. Further efforts are needed to explore the experimental conditions for efficient and specific gene editing in this cell type.

### ICE accuracy validation

We initially verified the sensitivity of ICE, TIDE and T7E1 analysis side by side. In artificial samples with incrementally increased INDELs frequencies from 1# to 3# edited cells, T7E1 assay failed to distinguish between them (Figure SA, C). In contrast, both ICE and TIDE revealed a clear upward trend (Figure S4 B, C). Next, we validated the accuracy of ICE analysis by comparing its predictions to actual edits obtained from real experiments targeting PLA2G6 gene (Figure S4D). The data showed that predicted total INDELs efficiency of 85% closely approached the observed 87% calculated from 50 sorted cells (Table S5). More importantly, the editing spectrum predicted by ICE highly matched the observed edits, accurately capturing the predominated editing types of +1, -1 and -11. Nevertheless, the data also revealed ICE failed to detect the edits occurring at frequency lower than 3% in the actual genotype of sorted cell clones (Table S5 and Figure S4E).

## Discussion

Despite more than a decade passing since the initial release of Cas9-sgRNA plasmids for gene editing in eukaryotic cells [25], Cas9-sgRNA plasmid-based approaches for constructing mutant cell lines remain prevalent worldwide. This might stem from their ease of use and versatility, allowing researchers to readily design and clone sgRNA sequences. While Cas9-sgRNA plasmids offers advantages in terms of accessibility, our study identified limitations associated with their use for generating mutant hPSCs lines. These limitations include potentially very low transfection efficiency and the inherent challenges of antibiotic selection. Antibiotic selection can be a complex process requiring careful optimization for each cell line, often involving a degree of trial-and-error to achieve the desired selection pressure. This process can significantly extend the timeline for generating a mutant hPSCs line, especially for newcomers to the field. Without experience in handling hPSCs, achieving a desired mutant line can take 2-3 months or even longer. These limitations highlight the need for a more efficient, user-friendly method for generating mutant hPSCs lines. The inducible Cas9 (iCas9) technique emerges as a promising solution. In previous hPSCs-iCas9 studies, the INDELs efficiencies achieved up to 68%, eliminating the need for laborious and variable antibiotic selection process. To fully harness its potential, we optimized several parameters with the goal of maximizing and stabilizing the INDELs introduction efficiency of hPSCs-iCas9.

In fact, most studies have overlooked the impact of cells’ nucleofection tolerance on editing efficiency. Our data revealed that H7 cells, being more vulnerable, consistently exhibit lower editing efficiency compared to the robust H9 cells. This discrepancy is most likely caused by H7 cells’ inability to withstand nucleofection stress, which leads to lower survival rates. This variation in tolerance might also partially explain the inconsistent editing efficiency reported in previous iCas9 studies using different cell lines [8,12,13]. However, our research offers a solution to address this inconsistency: repeated nucleofection. Though nucleofection frequency was initially not within the scope of our consideration, we unexpectedly discovered that repeating nucleofection effectively overcomes the hurdle of cells’ intolerance to nucleofection stress and boosts INDELs efficiency by up to threefold. In vulnerable H7 cells, for example, this approach increased INDELs from 36% achieved with the first nucleofection to 87% (ACE2-2# sgRNA, Figure 3D).

Previous hPSCs-iCas9 studies have commonly utilized plasmid-sgRNA (sgRNA delivered by plasmid transfection) or IVT-sgRNA, while the performance of CMS-sgRNA has been infrequently documented. Our data convincingly demonstrate that CMS-sgRNA consistently outperformed IVT-sgRNA in producing INDELs, achieving up to a threefold increase. This finding indicates that sgRNA stability is a critical factor influencing gene editing outcomes. Compared to plasmid-sgRNA, which are typically large (about 4000 nt), the compact size of CMS-sgRNA (around 150 nt) greatly favors transfections efficiency. Additionally, plasmid-sgRNA requires laborious molecular cloning and plasmid maxi-preparation work, while CMS-sgRNA can be readily synthesized by commercial vendors within a week. This translates to substantial time savings for researchers, especially in experiments requiring significant sgRNA quantities.

In addition to inducing INDELs at a single locus, the optimized hPSCs-iCas9 system achieved over 80% and 75% efficiency on each target locus in double and triple gene knockout experiments, respectively. This significantly surpasses previously reported efficiencies [12] and offers greater consistency compared to episomal plasmid approaches [26]. Furthermore, our optimized system excels at excising DNA fragments, achieving over 37% homozygous knockout for a 1.2 kb fragment removal. This efficiency rivals that of Cas9-sgRNA plasmid experiments, which however, necessitate extensive expertise in antibiotic selection [27]. Paired sgRNAs offers several key advantages over single sgRNA-induced small indels for knocking out genes. Firstly, it enables the removal of multiple or even all exons of a target gene, thereby maximizing the likelihood of eliminating the corresponding protein. Secondly, paired sgRNAs exhibits broad applicability, allowing for the deletion of genomic regulatory regions or non-coding RNAs [27]. This significantly expands the spectrum of genetic modifications achievable in hPSCs, facilitating more comprehensive studies of gene function and disease modeling.

We also investigated the optimized system’s competency for single-base substitutions by selecting a specific point mutation case to assess the maximum knock-in potential achievable. However, HDR-mediated knock-in efficiency only reached to average of 25%, showing limited improvement over previous studies [8]. Since numerous reports highlighting the competition between NHEJ and HDR pathways during Cas9-mediated editing [28,29], and considering the optimized system’s proficiency in NHEJ-mediated INDLEs, it may not be the preferred choice for HDR experiments. With the recent advancements in base editing and prime editing techniques, we recommend using these more precise tools for single-base substitutions instead of CRISPR/Cas9.

This study has been underscoring the critical role of sgRNA selection in gene knockout experiment and solidifying its position as an essential step in the refined workflow (Figure 7). Cleavage efficacy is a key criterion for sgRNA design as it has a direct impact on introducing INDELs. Although numerous free online scoring algorithms exist to aid researchers in sgRNA selection, identifying the most reliable one, especially when scores conflict, can be challenging. Our hPSCs-iCas9 system offers a unique advantage for sgRNA evaluation due to efficient RNA transfection into hPSCs and uniform Cas9 expression across the cell population upon Dox administration. Essentially, all Cas9-expressing cells are exposed to the sgRNA, providing an unbiased platform to assess sgRNA cleavage efficiency through real-world experiments. By comparing these experimental results with predicted scores, we found that the Benchling algorithm outperformed both CCTOP and Deephf in terms of consistency and accuracy. It is also important to note that Benchling has its limitations, as exemplified by its inability to distinguish slight differences between sgRNAs 1# and 2# targeting PHF19 (Figure S2E), where the predicted scores contradicted the experimental results.

**Figure 7:**
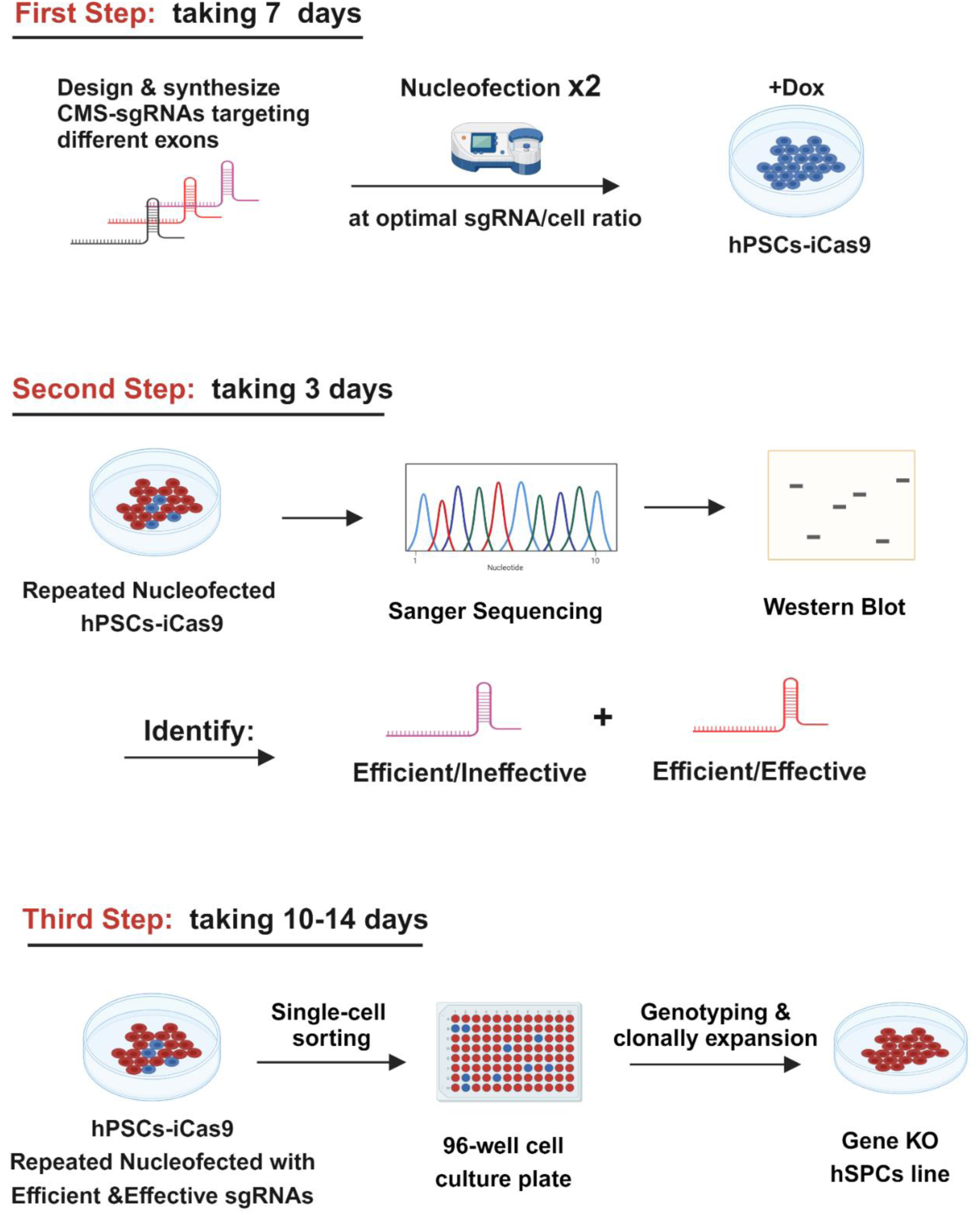
Refined workflow for establishing Gene knockout hPSCs lines. The refined workflow involves following steps: First step: (1) Design and order a set of CMS-sgRNAs targeting different exons of the target gene. (2) Add Dox to hPSC-iCas9 cells 12 hours prior to nucleofection experiment. (3) Conduct repeated nucleofection. typically completed within 7 days. Second step: Employ ICE to analyze Sanger sequencing results and conduct Western blot to detect target protein expression level. Based on these results, identify the efficient and effective sgRNAs. which takes approximately 3 days. Third step: (1) Perform single cell sorting for the edited cells pool nucleofected with efficient and effective sgRNA. (2) Culture sorted cells in 96-well plates and clonally expand them. (3) Genotype individual cell clone through Sanger sequencing to obtain the desired edited cell clones.

Due to exon skipping [30], nonsense-mediated decay (NMD) escape [31] and other unexplored mechanisms, NHEJ-mediated non-triplet INDELs in target genes may not always result in mRNA degradation and protein elimination. Our strategy leverages the high and stable INDELs efficiency achieved by the optimized hPSCs-iCas9 system and employs Western blot to detect target proteins in cells nucleofected with either effective or ineffective sgRNAs. This approach provides a highly practical method for determining sgRNA’s effectiveness by prioritizing functionality over understanding the exact mechanisms behind sgRNA ineffectiveness, which is not crucial for generating mutant hPSCs lines. The key benefit lies in identifying and discarding ineffective sgRNAs early on, preventing wasted effort in downstream experiments. However, this method has its limitation, as it cannot be applied to detect proteins expressed only in specialized cells, such as cardiac sarcomeric proteins, which are not expressed in hPSCs.

While next-generation sequencing (NGS) analysis of PCR amplicons (Amp-seq) is a standard method for quantifying INDELs and knock-in in gene editing experiments, its high cost and time requirements make it impractical for extensive parameter optimization studies. Therefore, identifying a low-cost tool for rapid assessment of optimization outcomes becomes crucial. This study compared three tools in analyzing INDELs: TIDE, its improved version ICE, and the enzyme-based T7E1 assay. Our data revealed limitations with the T7E1 assay, which is consistent with previously reported findings that T7E1 struggles to distinguish editing efficiencies above 30%, and its signal correlates more with indel distribution than overall sgRNA activity [32,33]. Therefore, T7E1 is better suited for confirming gene editing events rather than precise efficiency quantification. In contrast, decomposition algorithms like ICE and TIDE offer valuable and informative analysis of Sanger sequencing data. These two tools could help researchers identify the predominant INDELs types and distinguish between homozygous, heterozygous, or mixed editing states in subcloned cell lines. This capability is particularly useful for establishing mutant cell lines with specific edits. Previously, achieving this level of detail required laborious and expensive plasmid TA-cloning followed by Sanger sequencing. In this study, by validating ICE’s analysis directly against the actual genotypes of single-cell sorted cell clones, we revealed a close match between ICE’s analysis and the observed genotypes, accurately reflecting both INDELs distribution and efficiency. Although ICE missed some minor alterations, its speed and cost-effectiveness make it a valuable tool for optimization studies and future gene knockout experiments.

Overall, the extensive optimization of experimental parameters in this study has culminated in a highly efficient gene-editing hPSCs-iCas9 system. This system delivers exceptional gene knockout efficiency and offer practical solutions to the challenges associated with sgRNA selection. By integrating these advancements, we have streamlined the workflow for generating gene knockout hPSCs lines (Figure 7). This refined protocol is not only user-friendly, easy to master, and achievable within 20-24 days, but also maximizes the likelihood of obtaining the correct edited cell clones.

## Acknowledgements

We gratefully acknowledge funding support from the National Key Research and Development Program of China (2021YFC2701703, 2023YFA0915002), CAMS Innovation Fund for Medical Sciences (CIFMS) (2021-RC310-12, 2022-I2M-2-001 and 2023-I2M-1-003), National High-Level Hospital Clinical Research Funding (2022-GSP-GG-7), Non-profit Central Research Institute Fund of Chinese Academy of Medical Sciences (2019PT320026), Shenzhen Fundamental Research Program (JCYJ20220531091615034 and ZDSYS20200923172000001) and Shenzhen High-level Hospital Construction Fund.

## Author contributions

XJL and FL conceived and designed the experiments. JN, JHG and YQR performed the major experiments. PFH, MZ, ZHZ and YY established ACE2 and TAZ knockout hPSCs lines. JN, YQR and RB analyzed the data. XJL, JN and RB contributed to the writing of the manuscript.

## Competing interests

The authors declare that they have no known competing financial interests or personal relationships that could have appeared to influence the work reported in this paper.

---

**Figure S1.**
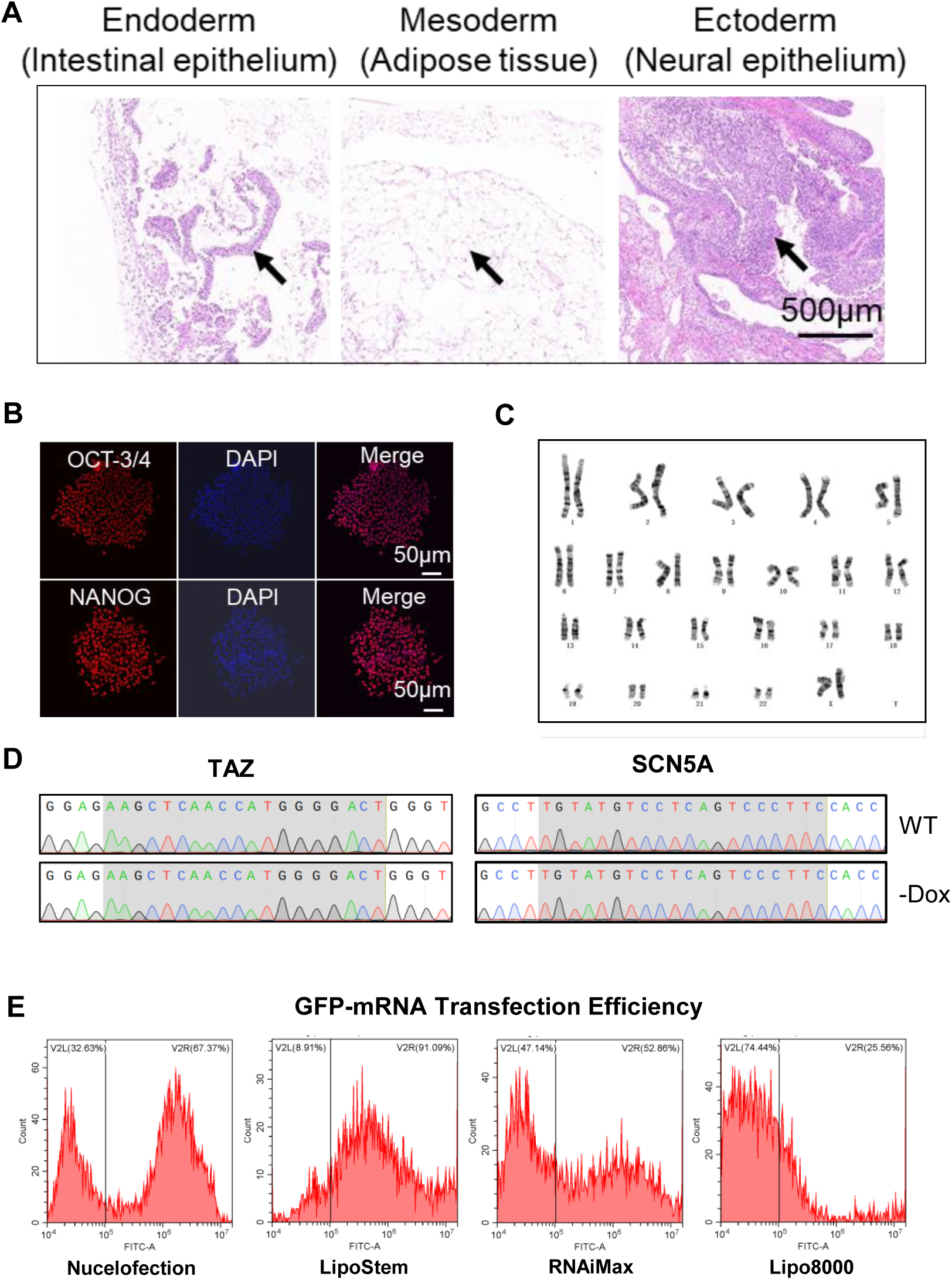
Characterization of hPSCs-iCas9. (A) Hematoxylin and eosin (HE) staining of the teratomas derived from H9-iCas9 cells. Arrows indicate the presence of endoderm (Intestinal epithelium), mesoderm (adipose), and ectoderm (neural epithelium). Scale bar=500 μm. (B) IFA staining for pluripotency markers OCT3/4 and NANOG in H9-iCas9. Cells were counterstained with DAPI to visualize nuclei. Scale bar=50 μm. (C) Karyotype analysis of H9-iCas9 line. (D) Sanger sequencing to detect the potential nuclease leakage. H9-iCas9 cells nucleofected with sgRNAs targeting the TAZ or SCN5A gene without Dox addition showed no visible edits in the suspected editing regions (areas in shadow) compared to Wild-type (WT). (E) Flow cytometry analysis of GFP-positive cells following GFP-mRNA transfection using different reagents. 8×10^5^ cells were transfected with 5 μg mRNA.

**Figure S2.**
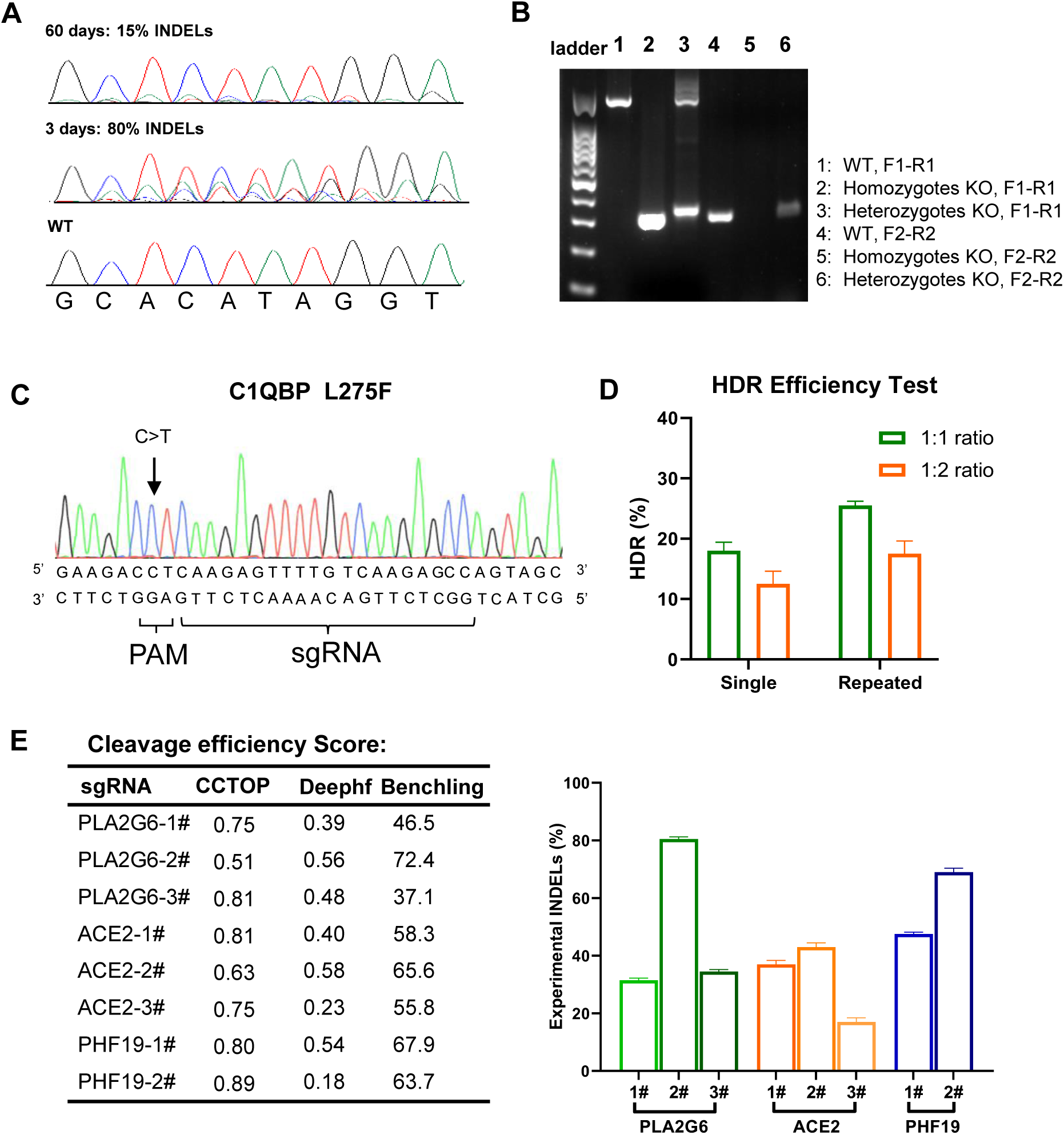
Gene editing performance of optimized hPSCs-iCas9. (A) Sanger sequencing chromatograms showing a notable reduction in INDELs incorporation from 3 to 60 days. (B) Representative gel electrophoresis of PCR products of paired sgRNA-mediated gene knockouts. Lane 1/2/3 depicts amplification with external primers (F1, R1), while lane 4/5/6 depicts amplification with internal primers (F2, R2). Lane 1/4 represents Wild-type, lane 2/5 represents homozygous knockout, and lane 3/6 represents heterozygous knockout. (C) Schematic diagram of the C1QBP-L275F (CTC>TTC) mutation and the sgRNA design for HDR. Following HDR, the mutation replaces the wild-type nucleic acid, thereby removing the PAM. (D) Optimization of sgRNA-to-ssODN ratio to enhance HDR efficiency. The total nucleic acid for nucleofection was fixed at 5 μg. Data represent the average of two replicates. (E) Validation of cleavage efficiency scores against the actual edits. Left: Cleavage efficiency predicted by three different scoring algorithms. Right: Experimental INDELs efficiency of sgRNAs targeting various gens determined in H9-iCas9. Data are shown as mean ± SD, n=3.

**Figure S3.**
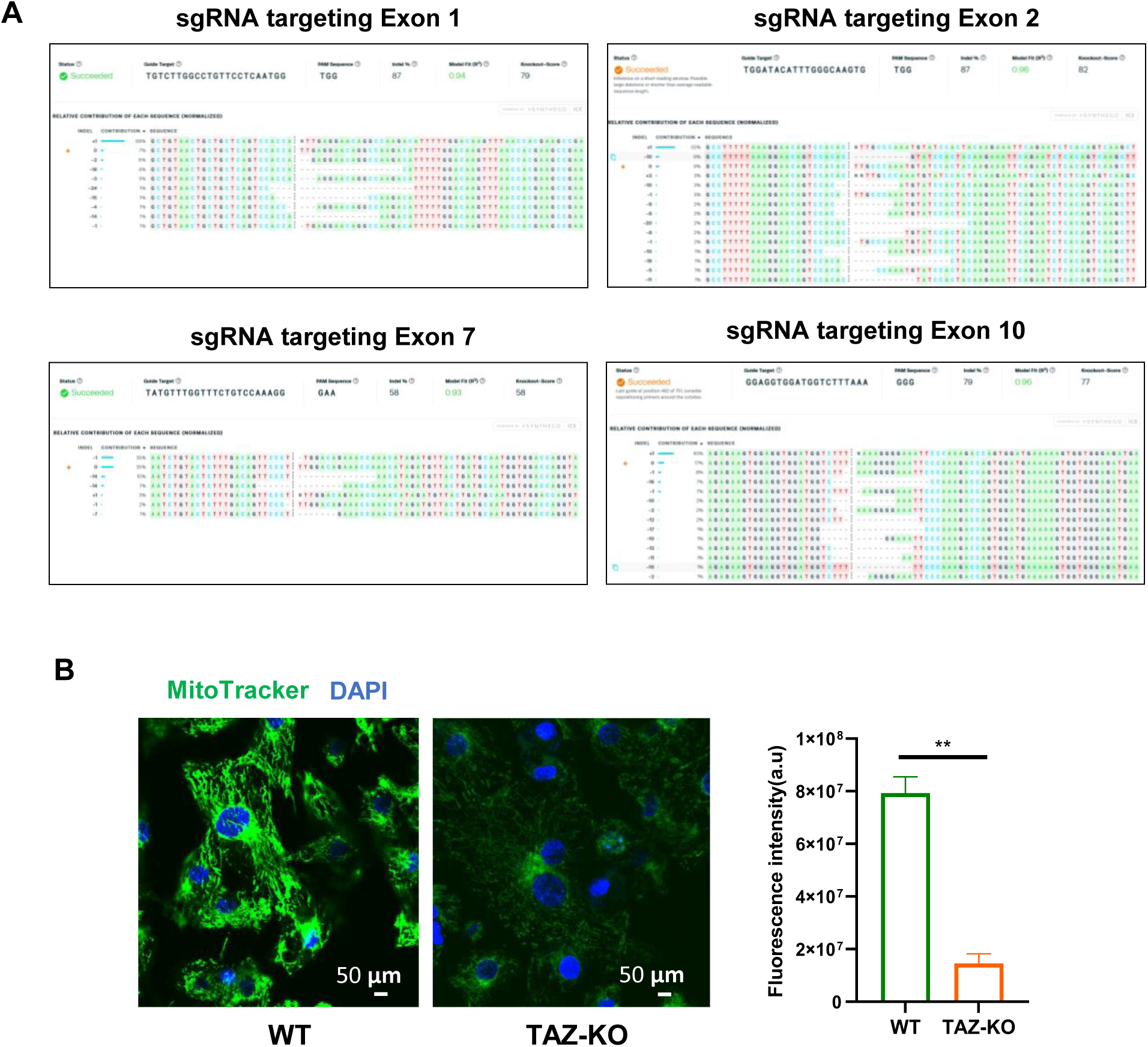
(A) ICE analysis of Sanger sequencing chromatograms for gene knockout on each exon. (B) Significantly reduced mitochondrial potential and activity of TAZ-KO cardiomyocytes. hPSCs-CMs were stained with MitoTracker Green and imaged with a confocal microscope. Left: Representative images, scale bar = 50 μm. Right: Quantitative analysis of mitochondrial staining intensity. Two-tailed *t*-test.

**Figure S4.**
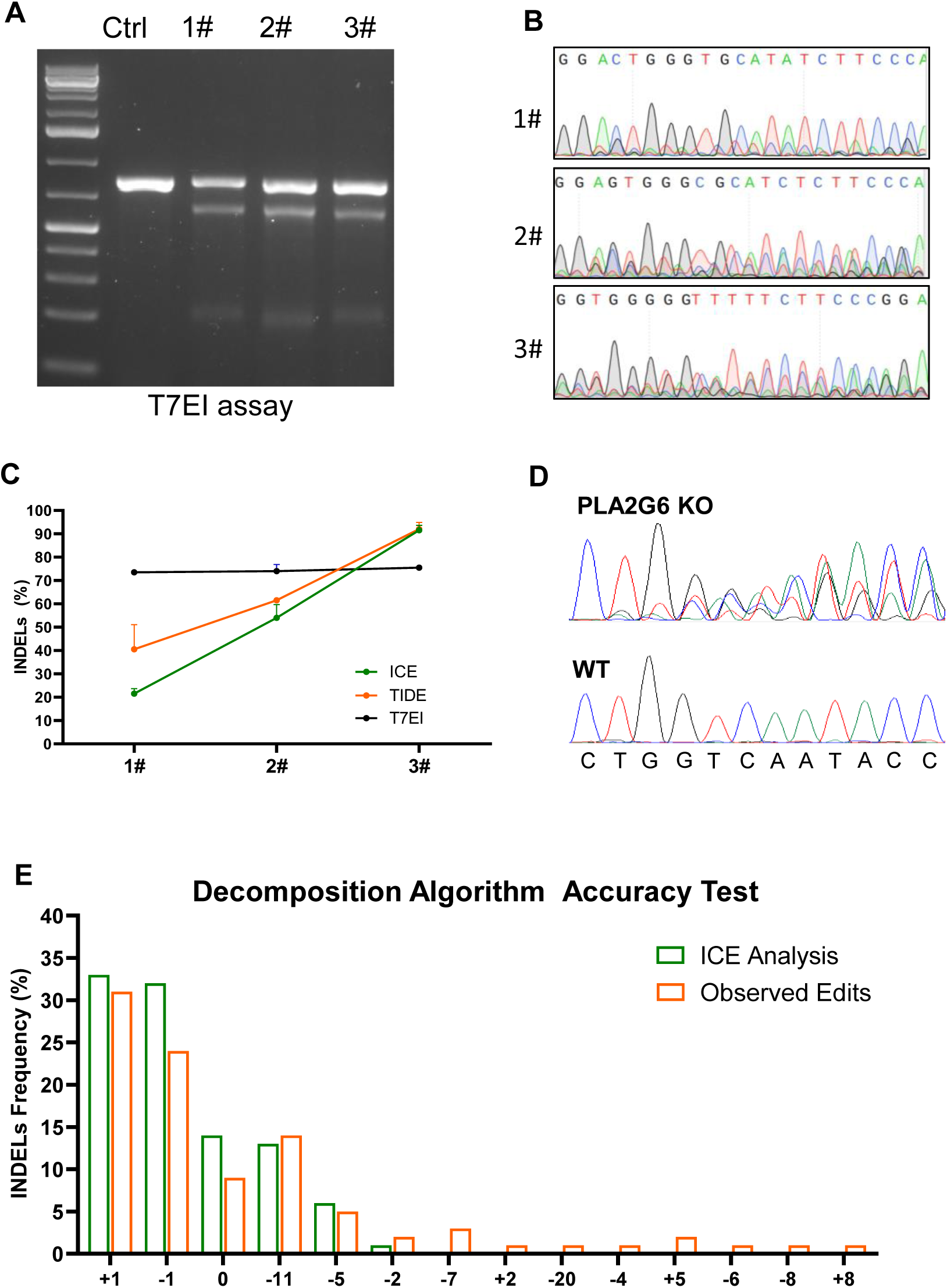
Validation of ICE analysis accuracy. (A) Representative gel electrophoresis image of PCR products from three edited cells pools (1# to 3#) digested by of 7TE1. (B) Sanger sequencing chromatograms of edited cells pools. (C) INDELs efficiency analyzed 7TE1, TIDE and ICE. n=3. (D) Sanger sequencing chromatograms of the edited cells pool nucleofected with sgRNA targeting PLA2G6. (E) Comparison of actual edits obtained from single-sorted cells with ICE-analyzed Sanger sequencing chromatograms.

## Notes

### Competing Interest Statement

The authors have declared no competing interest.

### Summary of Updates

the order of author list was corrected, as my name (Corresponding)should be at the last position.

