## Supplemental tables for "Optimization of gene knockout approaches and practical solutions to sgRNA selection challenges in hPSCs with inducible Cas9 system"

**Table S1.** PCR Primers used in this study.

| **Name** | **Sequence** | **Description** |
| --- | --- | --- |
| TAZ-T7EI-F | ATAGTCCCCAACACATGGGC | PCR for T7EI assays |
| TAZ-T7EI-R | TGCAGCACAGTGGACAGAAG | PCR for T7EI assays |
| TAZ-F | TAAGCTAACCTGTCACCCCA | PCR for INDELs sequencing |
| TAZ-R | AGAGCACAGAGGCGAGGCTT | PCR for INDELs sequencing |
| ACE2-1#-F | AGTCTAGGGAAAGTCATTCAGTGGATG | PCR for INDELs sequencing |
| ACE2-1#-R | CGGGCAGTAATCTAATCTTTAAGAGGCAA | PCR for INDELs sequencing |
| ACE2-2#-F | CCTTGACCTTCAGCGGAGT | PCR for INDELs sequencing |
| ACE2-2#-R | AAAATGTTCACAAACGTACCCGT | PCR for INDELs sequencing |
| ACE2-7#-F | GGGATTACAGGAGTACTCTTCATGAAGT | PCR for INDELs sequencing |
| ACE2-7#-R | GCTTTATTGGGTCCATGGGCT | PCR for INDELs sequencing |
| ACE2-10#-F | GACTGGGAAAGTGGATGAAATAGCTCT | PCR for INDELs sequencing |
| ACE2-10#-R | GTTTTTTGGGGTTGAGGACCATACC | PCR for INDELs sequencing |
| PCDH7-F | CAGTCGGTGGTGGAGGTTTAC | PCR for INDELs sequencing |
| PCDH7-R | AAAGCGAGACTGTAGCTGTGG | PCR for INDELs sequencing |
| C1QBP-F | TTGGACTGGGTGAGTGCTTG | PCR for INDELs &HDR sequencing |
| C1QBP-R | TCAAAGCACATGTCTAGCTTCAGA | PCR for INDELs & HDR sequencing |
| PHF19-F | ACCTGTTCTTCTGCTCCGTG | PCR for INDELs sequencing |
| PHF19-R | ATGCAGCAACTCAAGGGTGA | PCR for INDELs sequencing |
| PHF19-F1 | GGTTGCCGAAGCTCATACCT | PCR for gene fragment deletion detection (Figure 5B) |
| PHF19-R1 | ATGCAGCAACTCAAGGGTGA | PCR for gene fragment deletion detection (Figure 5B) |
| PHF19-F2 | AACCGACGGAGCAAAGCTAA | PCR for gene segment deletion detection (Figure 5B) |
| PHF19-R2 | TTCCCCACCTACCTTCCCAT | PCR for gene fragment deletion detection (Figure 5B) |
| SCN5A-F | AGATCTCCCTCACCTCCACC | PCR for INDELs sequencing |
| SCN5A-R | GCCAATATCAGAGCCCGACA | PCR for INDELs sequencing |
| PLA2G6-F | TGATCTGGGTGTCTGTGCAGGAAA | PCR for INDELs sequencing (Figure 3 D) |
| PLA2G6-R | GCTTCTCGGCCAATAAGACCTCC | PCR for INDELs sequencing (Figure 3 D) |
| PLA2G6-1#-F | AGTAGGAGGAAGTAGAAGTGCTGAGTAAG | PCR for INDELs sequencing (Figure 5B) |
| PLA2G6-1#-R | GGGAGTCACATAAGGGTTAATGGAGGAC | PCR for INDELs sequencing  (Figure 5B) |
| PLA2G6-2#-F | TGATCTGGGTGTCTGTGCAGGAAA | PCR for INDELs sequencing  (Figure 5B) |
| PLA2G6-2#-R | GCTTCTCGGCCAATAAGACCTCC | PCR for INDELs sequencing  (Figure 5B) |
| PLA2G6-3#-F | ACCCACCTGACCACAGCATG | PCR for INDELs sequencing  (Figure 5B) |
| PLA2G6-3#-R | GGGTGCAGAGAGTAAAGCCCTG | PCR for INDELs sequencing  (Figure 5B) |

**Table S2.** Antibodies used in this study.

| **Antibody** | **Source** | **Cat. No.** | **Dilution** |
| --- | --- | --- | --- |
| Cas9(7A9-3A3) Mouse mAb(Alexa Fluor® 647 Conjugate) | Cell Signaling Technology | 48796S | FACS 1:50 |
| Cas9(7A9-3A3) | Santa Cruz Biotechnology | SC-517386 | WB: 1:500 |
| GAPDH | Santa Cruz | SC-47724 | WB:1:500 |
| OCT3/4 | Santa Cruz | SC-5279 | IFA:1:200 |
| NANOG | Santa Cruz | SC-293121 | IFA:1:100 |
| cTNT | Santa Cruz | SC-20025 | IFA:1:50 WB:1:200 |
| ACE2 | invitrogen | MA5-32307 | WB:1:1000 |
| DAPI | Thermo Fisher | 62248 | IFA: 1:1000 |
| Anti-Mouse IgG (Alexa Fluor® 647 Conjugate) | Cell Signaling Technology | 4410 | FACS:1:1000 |
| Goat anti-Rabbit HRP | Absin | abs20002 | WB:1:5000 |
| Goat anti-Mouse HRP | Absin | abs20001 | WB:1:5000 |

**Table S3.** sgRNA and ssODN used in this study.

| **Name** | **Sequence** | **Description** |
| --- | --- | --- |
| TAZ sgRNA | GAAGCTCAACCATGGGGACT | INDELs efficiency test |
| SCN5A sgRNA | GAAGGGACTGAGGACATACA | INDELs efficiency test |
| PHF19 sgRNA | TCTGTACCCCCAGATTATAG | INDELs efficiency test |
| PCDH7 sgRNA | TTTCTATTGAAAATGACACG | INDELs efficiency test |
| PLA2G6 sgRNA | CACTGAAGGTATTGACCAGG | INDELs efficiency test |
| PHF19 sgRNA-1 | GTGCAGGCAGTGGTTCCACG | Gene fragment deletion efficiency test |
| PHF19 sgRNA-2 | TCTGTACCCCCAGATTATAG | Gene fragment deletion efficiency test |
| CIQBP sgRNA | GGCTCTTGACAAAACTCTTG | HDR efficiency test |
| TAZ-1# sgRNA | TTTTGGAGAAGCTCAACCAT | cleavage efficiency test |
| TAZ-2# sgRNA | TTTGGAGAAGCTCAACCATG | cleavage efficiency test |
| TAZ-3# sgRNA | ATTTTGGAGAAGCTCAACCA | cleavage efficiency test |
| TAZ-4# sgRNA | GAAGGGGATGGACTTCATTT | cleavage efficiency test |
| TAZ-5# sgRNA | AGATATGCACCCAGTCCCCA | cleavage efficiency test |
| PLA2G6-1# sgRNA | GGTCCGTTCCCCAACTTCCT | cleavage efficiency test |
| PLA2G6-2# sgRNA | CACTGAAGGTATTGACCAGG | cleavage efficiency test |
| PLA2G6-3# sgRNA | CCAGGATGAACGCTGGCTTC | cleavage efficiency test |
| ACE2 -1# sgRNA | TGTCTTGGCCTGTTCCTCAATGG | cleavage efficiency test |
| ACE2 -2# sgRNA | TGGATACATTTGGGCAAGTG | cleavage efficiency test |
| ACE2-7# sgRNA | TATGTTTGGTTTCTGTCCAAAGG | cleavage efficiency test |
| ACE2-10# sgRNA | GGAGGTGGATGGTCTTTAAA | cleavage efficiency test |
| CIQBP ssODN | AGCTCAGCACAGCCCTGGAGCACCAGGAGTACATTACTTTTCTTGAAGACTTCAAGAGTTTTGTCAAGAGCCAGTAGAGCAGACAGATGCTGAAAGCCATA | homology directed repair  template |

**Table S4. Off-target Sequencing for TAZ sgRNA**

Targeted sequencing did not detect off-target editing by the TAZ gRNA. The TAZ-targeted and top 8 off-target sites predicted by CCTop were PCR amplified and evaluated by Sanger sequencing. Off target sites are shown with mismatches to gRNA sequence shown in red. Check indicates the wild-type sequence.

| **Coordinates** | **Gene** | **Sequence** | **Wild-type** | **TAZ** |
| --- | --- | --- | --- | --- |
| chrX:  154419583-154419605 | TAZ | GAAGCTCAACCATGGGGACT | √ | Mutation |
| chr1:  237784936-237784958 | RYR2 | GCACCTCAGGCATGGGGACT | √ | √ |
| chr3:  10127061-10127083 | BRK1 | AAAGTCCAAACATGGGGACT | √ | √ |
| chr7:  36096500-36096522 | RP11-  196O2.1 | GTGGCACAGCCATGGGGACT | √ | √ |
| chr5:  7924664-7924686 | CTD-  2072I24.1 | CCAGCACAAGCATGGGGACT | √ | √ |
| chr11:  124882759-124882781 | RP11-  664I21.5 | GAGGCAGGACCATGGGGACT | √ | √ |
| chr7:  38606132-38606154 | AMPH | AAACCTAAAGCATGGGGACT | √ | √ |
| chr7:  27899919-27899941 | JAZF1 | GGAGCTGGAGCATGGGGACT | √ | √ |
| chr4:  117898307-117898329 | AC108056.1 | GAAGCCCCACCTTGGGGACT | √ | √ |

**Table S5. ICE predication versus observed edits**

| **Edit types (bp)** | **ICE predicated** | **Observed** |
| --- | --- | --- |
| +1 | 33% | 31% |
| -1 | 32% | 24% |
| 0 | 14% | 9% |
| -11 | 13% | 14% |
| -5 | 6% | 5% |
| -2 | 1% | 2% |
| -7 | 0 | 3% |
| +2 | 0 | 1% |
| -20 | 0 | 1% |
| -4 | 0 | 1% |
| +5 | 0 | 2% |
| -6 | 0 | 1% |
| -8 | 0 | 1% |
| +8 | 0 | 1% |
| Total INDELs | 85% | 87% |
